# Genome size changes by duplication, insertion and divergence in Caenorhabditis worms

**DOI:** 10.1101/2022.05.20.492698

**Authors:** Joshua D. Millwood, John M. Sutton, Paula E. Adams, Jason Pienaar, Janna L. Fierst

## Abstract

Genome size has been measurable since the 1940s but we still do not understand the basis of genome size variation. *Caenorhabditis* nematodes show strong conservation of chromosome number but vary in genome size between closely related species. Androdioecy, where populations are composed of males and self-fertile hermaphrodites, has evolved from outcrossing, female-male dioecy, three times in this group. Androdioecious genomes are 10-30% smaller than dioecious species but large phylogenetic distances and rapid protein evolution have made it difficult to pinpoint the basis of these changes. Here, we analyze the genome sequences of *Caenorhabditis* and and test three hypotheses explaining genome evolution: 1) genomes evolve through deletions and ‘genome shrinkage’ in androdioecious species; 2) genome size is determined by transposable element (TE) expansion and DNA loss through large deletions (the ‘accordion model’); and 3) TE dynamics differ in androdioecious and dioecious species. We find no evidence for these hypotheses in *Caenorhabditis*. Across both short and long evolutionary distances *Caenorhabditis* genomes evolve through small structural variant (SV) mutations including frequent duplications and insertions, predominantly in genic regions. *Caenorhabditis* have rapid rates of gene family expansion and contraction and we identify 71 protein families with significant, parallel decreases across self-fertile *Caenorhabditis*. These include genes involved in the sensory system, regulatory proteins and membrane-associated immune responses, reflecting the shifting selection pressures that result from self-fertility. Our results suggest that the rules governing genome evolution differ between organisms based on ecology, life style and reproductive system.

## Introduction

Genetic variation is the fuel for adaptation (Fisher, 1930). Molecular evolutionary studies have focused on genomic variation at single nucleotide polymorphisms (SNPs) but gene deletions, duplications and other SV mutations may play important roles in adaptation and species divergence. Long-read DNA sequencing technologies have recently expanded our capability to accurately characterize SVs and we can now address the evolutionary significance of SVs in genomic evolution.

Nematodes have high rates of genomic polymorphisms and are a compelling system for studying the evolution of genetic and genomic variation. Outcrossing dioecious *Caenorhabditis* are characterized by molecular ‘hyper diversity’ and little linkage disequilibrium (Dey *et al*., 2013; Cutter *et al*., 2006) while the self-fertile androdioecious *C. elegans* has remarkably low levels of genetic variation (Cutter *et al*., 2009) and little global population structure (Andersen *et al*., 2012). Despite this there is substantial genomic divergence between *C. elegans* strains. The genome of the ‘Hawaiian’ CB4856 strain contains an extra 4Mb of genomic sequence when compared with the laboratory standard ‘Bristol’ N2 (Kim *et al*., 2019; Thompson *et al*., 2015). Sequencing and analysis across hundreds of wild-collected *C. elegans* strains indicates that 2-10% difference in genome size is common (Cook *et al*., 2017) with copy number (Maydan *et al*., 2010) and gene presence-absence variation (Lee *et al*., 2022) contributing to these differences.

Across the *Caenorhabditis* group self-fertile species have genomes 10-30% smaller than related outcrossing species (Bird *et al*., 2005; Haag *et al*., 2007). Similar patterns of smaller self-fertile genomes and larger outcrossing genomes have been reported in plants including *Arabidopsis* (Hu *et al*., 2011) and *Capsella* (Slotte *et al*., 2013) with equivocal support for different mechanisms of genome expansion and reduction. In *Caenorhabditis* genome size differences reflect outcrossing species higher gene number and protein-coding genome content (Stevens *et al*., 2019; Teterina *et al*., 2020; Thomas *et al*., 2012; Fierst *et al*., 2015; Noble *et al*., 2021), but it is unclear if this divergence occurred through ‘shrinkage’ across self-fertile species (Yin *et al*., 2018) or growth across outcrossing species (Kanzaki *et al*., 2018). There is also a substantial phylogenetic component to genome size in *Caenorhabditis*, further complicating analyses (Stevens *et al*., 2019).

Mutation accumulation studies focusing on the self-fertile *C. elegans* show SV mutations are common. Gene deletion and duplication are frequent (Lipinski *et al*., 2011; Farslow *et al*., 2015; Katju and Bergthorsson, 2013) with duplications occuring at an estimated 2.9*x*10^*−*5^*/*gene per generation and deletions at 5*x*10^*−*6^*/*gene per generation (Konrad *et al*., 2018). In comparison, single nucleotide base substitutions occur at 10^*−*9^ to 10^*−*8^ per generation (Denver *et al*., 2004).

The evolutionary significance of these SV patterns has been difficult to address in *C. elegans* because the most closely related known species, *C. inopinata*, diverged roughly 12 million years ago (Kanzaki *et al*., 2018). Outcrossing species have resisted the inbreeding necessary to create homozygous genome sequences (Dolgin *et al*., 2007) and residual allelism has prevented rigorous genomic analyses (Barriere *et al*., 2009). Self-fertility has evolved at least three times in the *Caenorhabditis* group but the known self-fertile species diverged more than 30 million years ago (Cutter, 2008). This span of divergence means that genomic comparisons between these species can not increase our understanding of SVs across evolutionary scales.

Here, we study SVs across a range of evolutionary divergence times. We start very small and compare the genomes of three strains of the outcrossing nematode *C. remanei*. We include *C. latens*, a dioecious species originally classified as a strain of *C. remanei* and later defined as a separate species (Dey *et al*., 2012). The *Caenorhabditis Elegans* group represents over 100 million years of evolutionary divergence and we broaden our comparisons to the outcrossing species *C. nigoni, C. inopinata* and *C. sinica* and the self-fertile *C. elegans, C. briggsae* and *C. tropicalis*.

Assembled genome sequences are available for more than 25 *Caenorhabditis* species but many of these remain fractured and incomplete (Stevens *et al*., 2019). The genic regions of these draft sequences are sufficient for phylogenetic reconstruction and other targeted analyses but not sufficient for fine-scale studies of genomic features. Transposable elements (TEs) retain high sequence similarity and these nearly-similar sequences are often collapsed or missing from draft assembled sequences (Ekblom and Wolf, 2014). Models of genome evolution often invoke TE-related changes and accurate quantification and characterization is necessary to test theoretical predictions and infer relationships between pattern and process. For example, the ‘accordion model’ of genome evolution proposes that genome size changes primarily through TE expansion and large segmental deletions (Kapusta *et al*., 2017). Population genetic theory predicts that Class I TEs involving an extrachromosomal RNA intermediate (retroelements) may be differentially affected by the evolution of self-fertility when compared with Class II TEs (DNA elements) with a ‘cut-and-paste’ mechanism (Boutin *et al*., 2012; Dolgin and Charlesworth, 2006). Here, we analyze chromosome-scale genome sequences that permit reliable characterization of protein-coding gene content, TE content and TE type. We focus on the *Caenorhabditis Elegans* group and test three hypotheses of genome evolution: 1) ‘genome shrinkage’ in self-fertile species (Yin *et al*., 2018); 2) the ‘accordion model’ (Kapusta *et al*., 2017); and 3) differing Class I/Class II TE dynamics in androdioecious and dioecious species (Boutin *et al*., 2012; Dolgin and Charlesworth, 2006).

We find *Caenorhabditis* genome evolution is characterized by numerous small-scale rearrangements. These occur frequently, even between closely related strains within species and the majority of rearrangements are within genes. TE-associated rearrangements are less frequent but have a larger mean and median size. Ancestral genome reconstructions show that duplicated sequence is the most common mode of genomic change with the larger outcrossing genomes due to an excess of duplications in comparison with the smaller self-fertile genomes. We find no evidence for the ‘accordion model’ of genome evolution in these worms, and no evidence for differential evolution of Class I and Class II TEs in outcrossing and self-fertile *Caenorhabditis*. Gene families show high birth and death rates across *Caenorhabditis* and there are no gene families changing in parallel across outcrossing *Caenorhabditis*. We find 71 gene families have decreased across self-fertile *Caenorhabditis* while no gene families have increased in parallel across these species. Significant, parallel reductions have occurred in self-fertile species sensory systems, regulatory proteins and membrane-associated immune responses. These protein changes reflect the shifts in selection that occur with self-fertility including a reduced need to find a mate or deal with pathogens. Overall, our results point to variable, dynamic rules of genome evolution across phylogenetic groups.

## Results

### Variation in protein-coding gene number between closely related Caenorhabditis

Similar to previous work (Thomas *et al*., 2012; Yin *et al*., 2018; Fierst *et al*., 2015) we found that genome size varied with reproductive mode (Fig. 1). The genomes of outcrossing *Caenorhabditis* were 120.37-130.48Mb in size while self-fertile *Caenorhabditis* genomes were 80.98-105.42 Mb. The number of protein-coding genes in these species also varied with reproductive mode with outcrossing genomes containing 21,443-34,696 genes while self-fertile genomes contained 19,997-21,210 genes. The fragmented *C. sinica* sequence (Supplementary Table 1) may have an artificially inflated estimated gene number of 34,696. Excluding *C. sinica* the mean number of genes in outcrossing genomes was 25,258 as compared to 20,714 in self-fertile genomes.

**Fig. 1:**
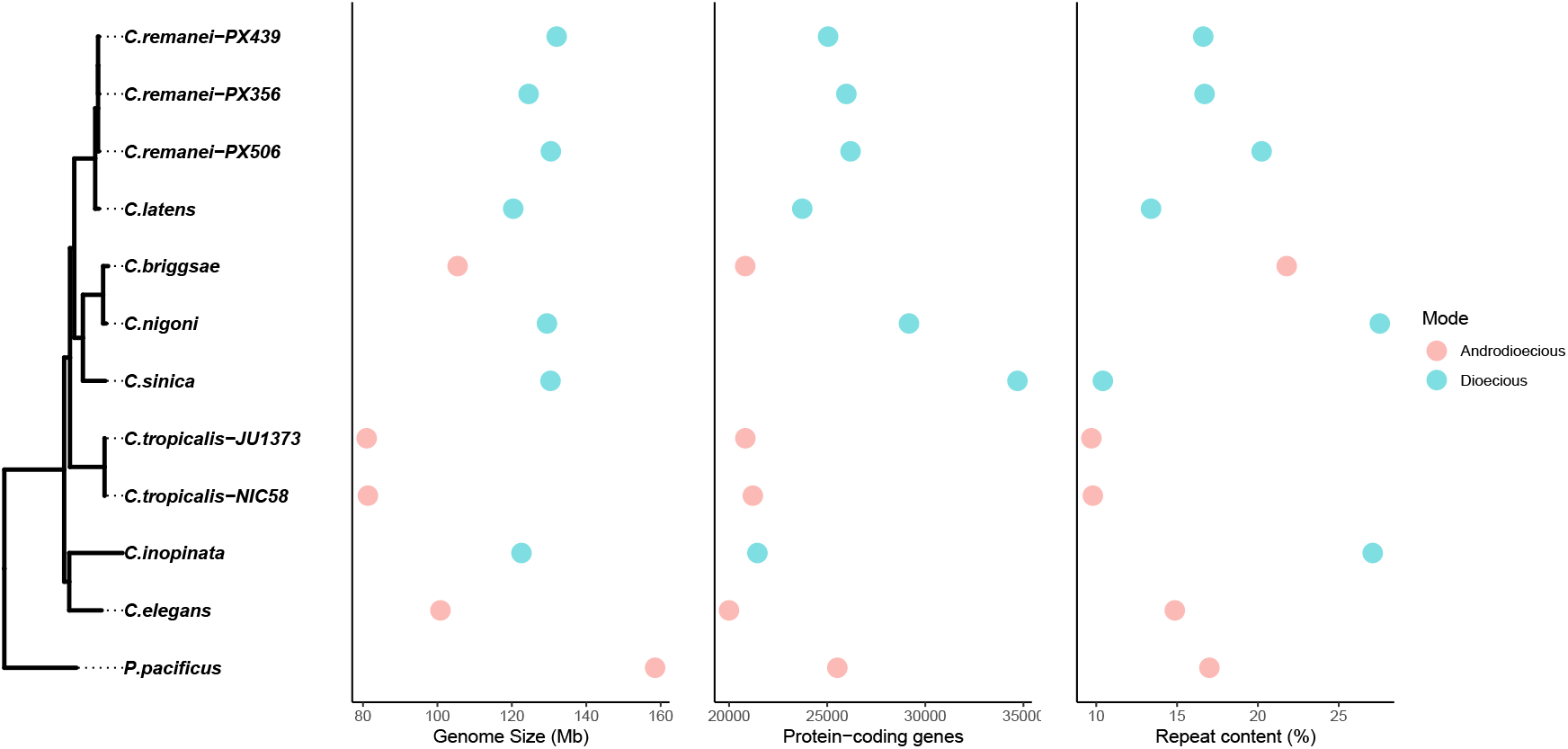
Genome size and number of protein-coding genes varied with reproductive mode while repeat content did not.

### Caenorhabditis genome evolution is characterized by extensive gene-associated insertions and duplications

Genomic changes from ancestor to child can occur in multiple ways (Fig. 2). The most common nucleotide changes across the *Caenorhabditis Elegans* group were insertions and duplications (Fig. 3). Deletions represented a small fraction of observed changes between both closely and more distantly related species pairs. Rapid nucleotide divergence could account for some of these differences as highly divergent nucleotide regions could escape classification as substitutions and result in an overrepresentation of ‘inserted’ sequences.

**Fig. 2:**
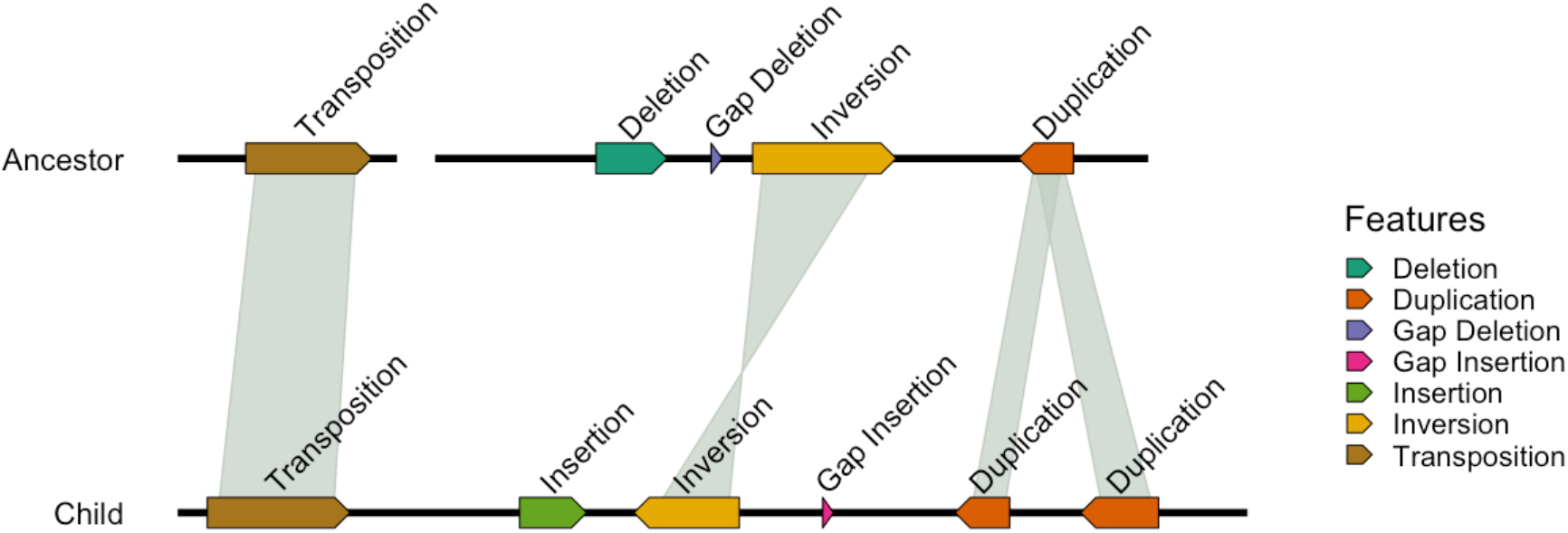
Genomic changes from ancestor to child can happen by transposition, small (gap) and large deletions, small (gap) and large insertions, inversions and duplications.

**Fig. 3:**
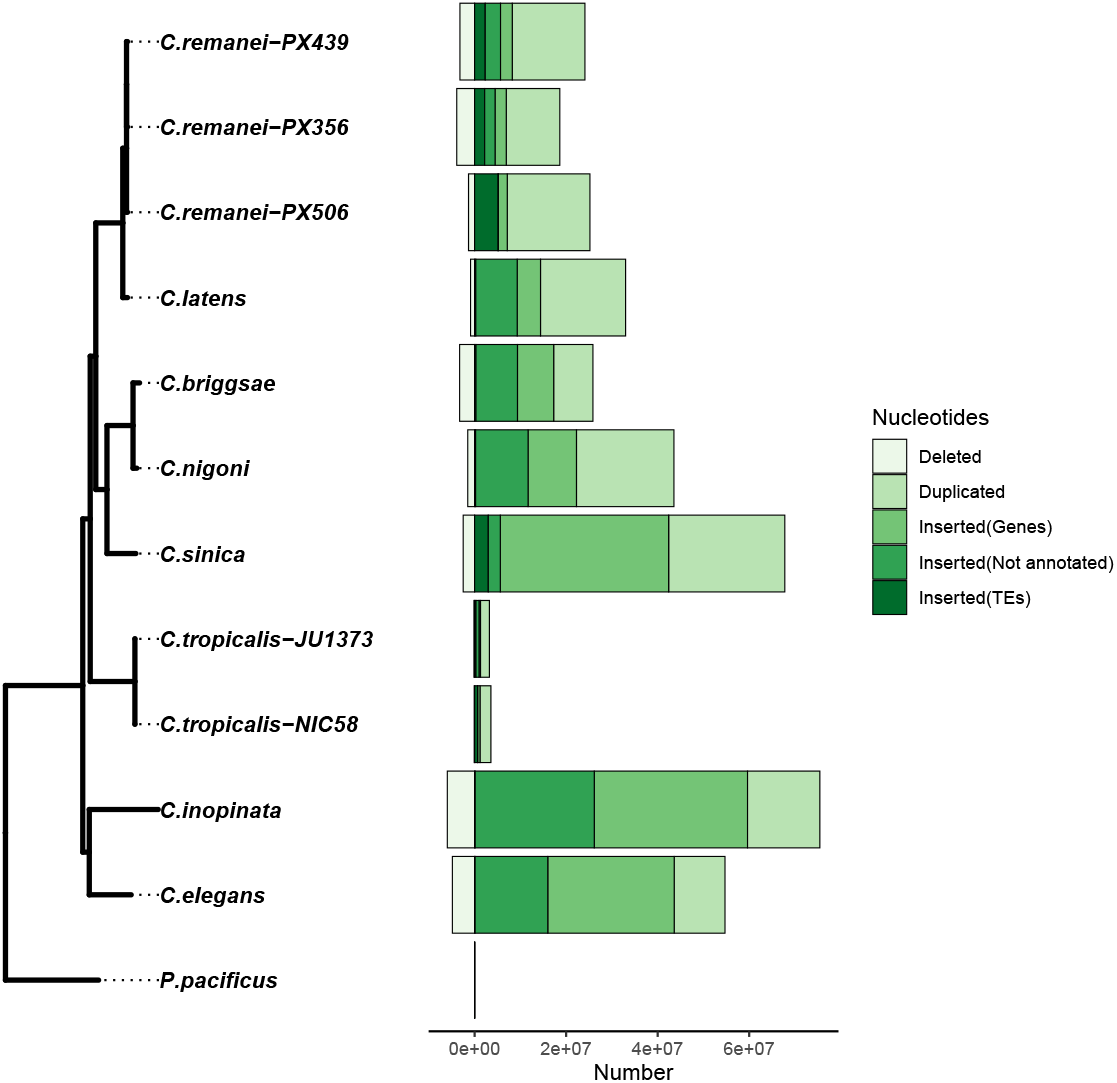
*Caenorhabditis* genome evolution is characterized by duplications and insertions with infrequent deletions. For most species insertions map to gene regions with the exception of *C. remanei* PX506.

For closely related species like the *C. remanei* /*C. latens* species group nucleotide changes can be tracked at a higher resolution. For example, for *C. remanei* PX506 we find that 1.33Mb of sequence was deleted relative to the ancestor (1% of the *C. remanei* PX506 genome size), 2% of nucleotides changed through substitution, 5% of the nucleotides were inserted and 14% of the nucleotides were duplicated (Supplementary Data 2). In comparison, 92% of the nucleotides were aligned/matched to the reconstructed ancestor sequence. Nucleotide changes can be encased in larger genomic rearrangements and multiply represented in the dataset and the total nucleotide changes sums to a representation greater than the current genome size (*i*.*e*., greater than 100%). Despite this, we can compare the relative contribution of each type of genomic change to genome size in the dataset and reject deletions as the predominant mode of genome evolution in this group (*df* = 21, *t* = 4.75, *p* = 5.47*x*10^*−*5^).

We associated the mutations identified by Progressive Cactus with genomic features in each dataset. We found that most genomic changes in the *Caenorhabditis Elegans* group occurred within genic sequences or unannotated regions (Fig. 3; *df* = 10, *t* = 2.46, *p* = 0.017). Changes within and close to TEs were less frequent but of larger mean and median size in all genomes. Inversions and transpositions occurred at a frequency of 1-2 orders of magnitude less than insertions with mean and median sizes 1-2 orders of magnitude greater than insertion-associated changes. The landscape of insertions, inversions and transpositions by chromosome is shown for the self-fertile *C. briggsae* in Fig. 4 and its outcrossing relative *C. nigoni* in Fig. 5. Both species showed similar patterns with mutations more frequent at the edges of chromosomes and slightly less frequent in chromosome centers.

**Fig. 4:**
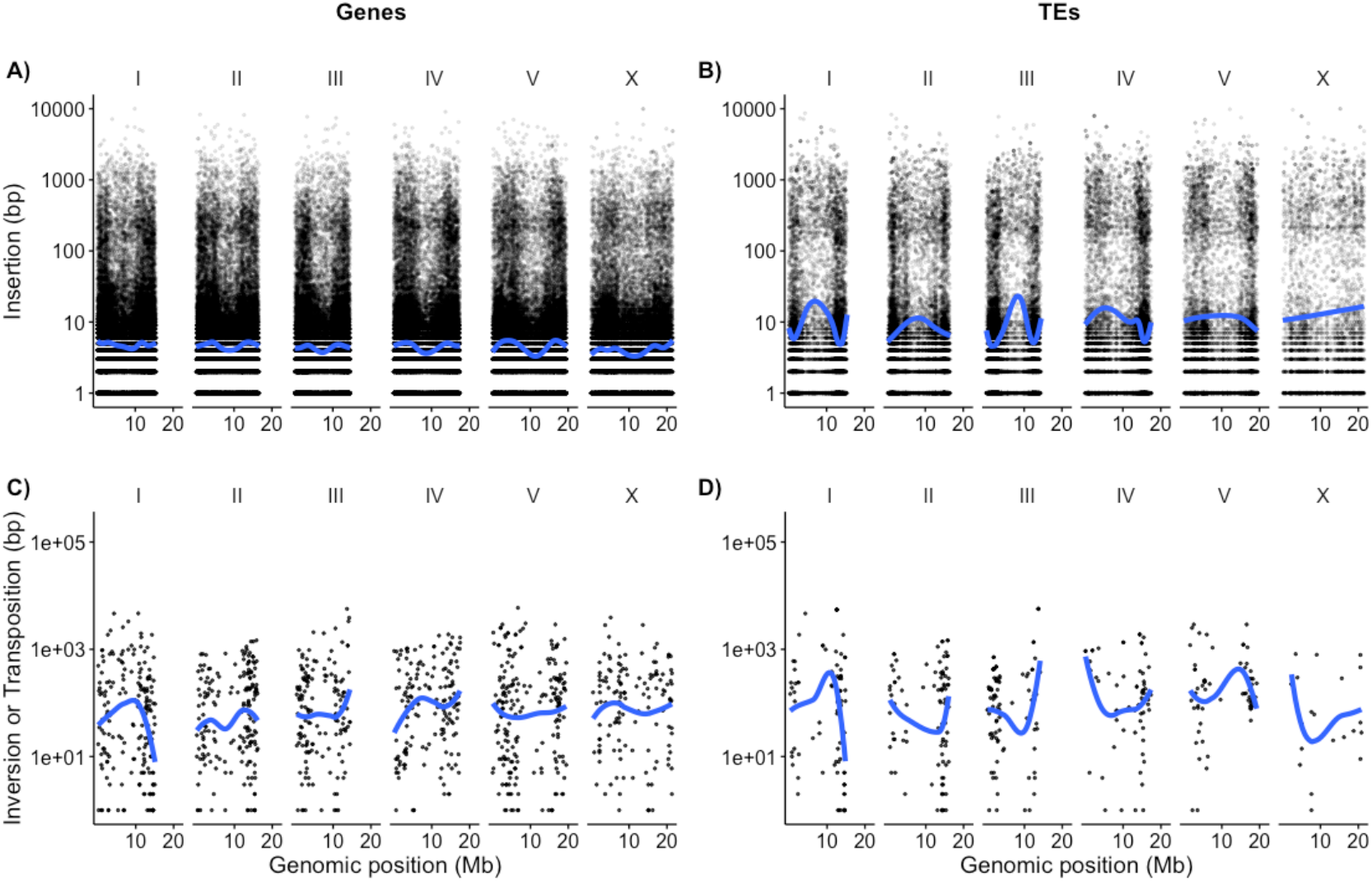
The most frequent type of genomic changes between *C. briggsae* and its ancestor were small gene-associated insertions.

**Fig. 5:**
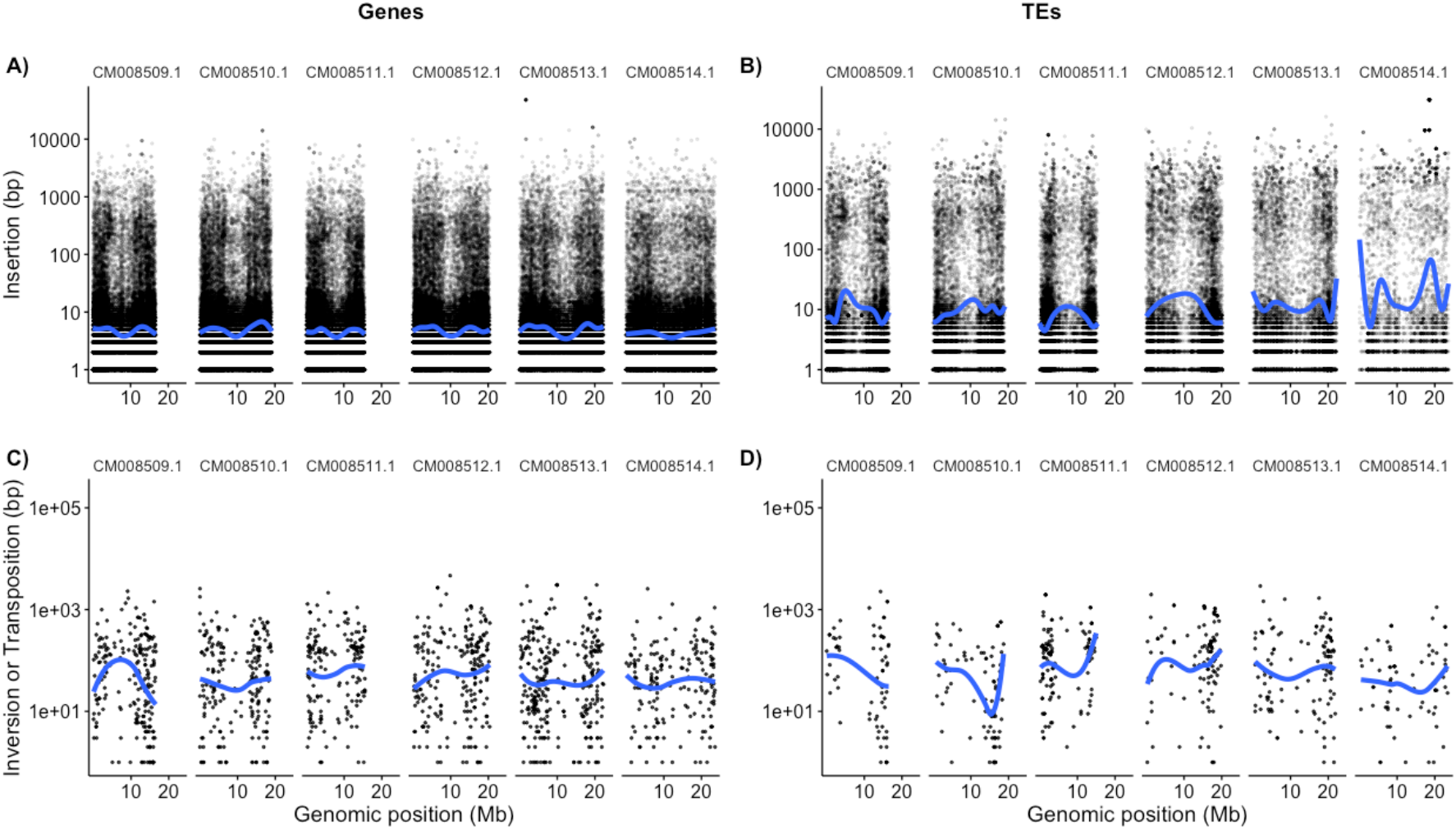
The genome of *C. nigoni*, the most closely related outcrossing species to *C. briggsae*, also diverged from their common ancestor primarily through small gene-associated insertions.

For the majority of the *Caenorhabditis* species mutations were distributed across all chromosomes equally with the exception of the *C. remanei* /*C. latens* species complex. For example, in *C. remanei* PX506 both gene-associated and TE-associated mutations were less frequent on the X chromosome (Fig. 6).

**Fig. 6:**
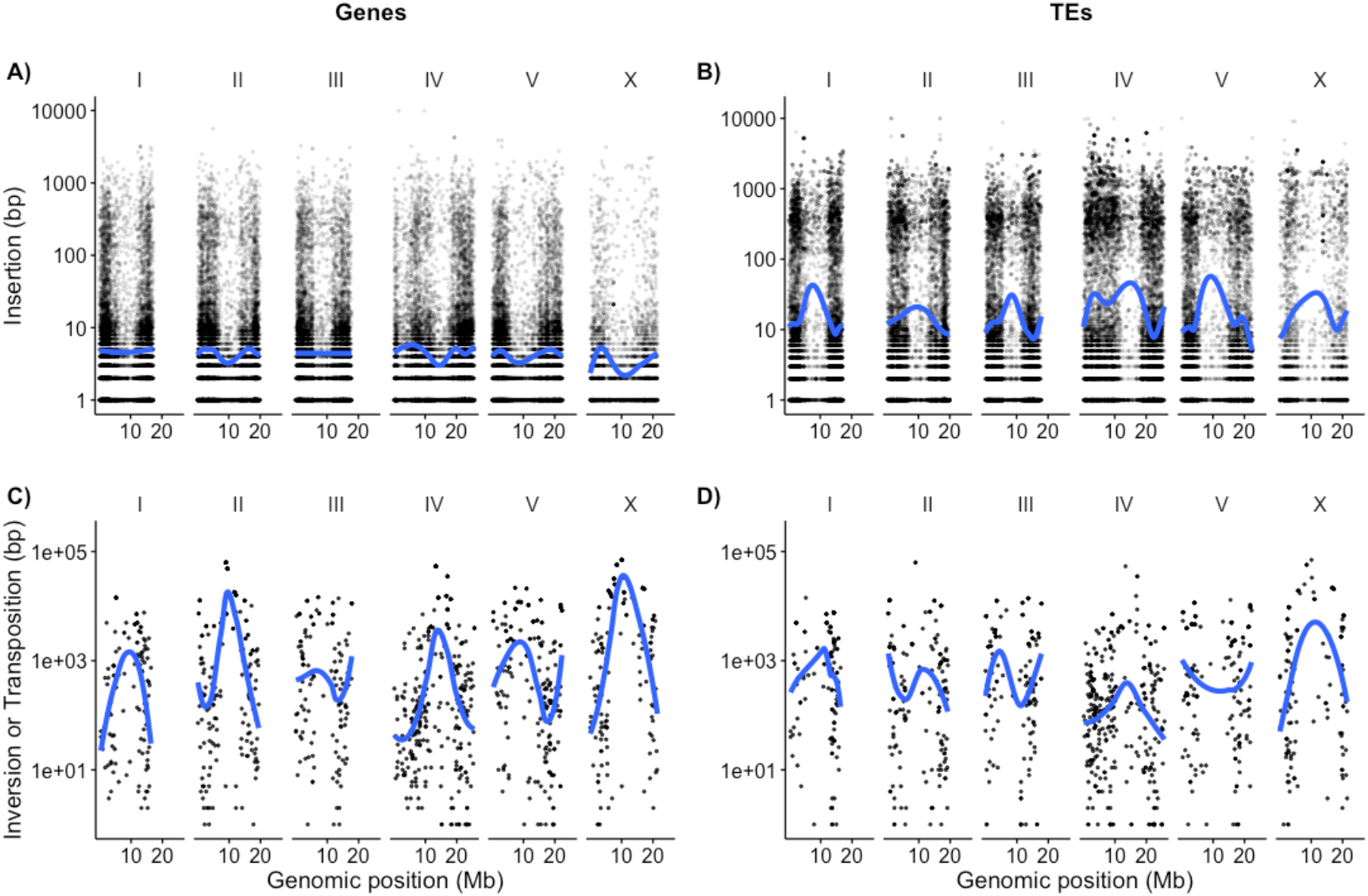
The outcrossing *C. remanei* PX506 diverged from its ancestor through small gene-associated insertions.

Insertions, inversions and tranpositions were calculated in the species genome coordinates while deletions and duplications were calculated in the ancestral genome coordinates. This limited our ability to infer overlap with genomic features as the ancestral genomes are not reconstructed at a sufficiently high resolution to annotate protein-coding genes and TEs. Duplications were more frequent than deletions and had larger mean and median sizes. Deletion and duplication patterns were similar for both self-fertile and outcrossing *Caenorhabditis*. For example, the mean and median sizes of deletions and duplications were similar for both the self-fertile *C. briggsae* and the outcrossing *C. nigoni* (Fig. 7).

**Fig. 7:**
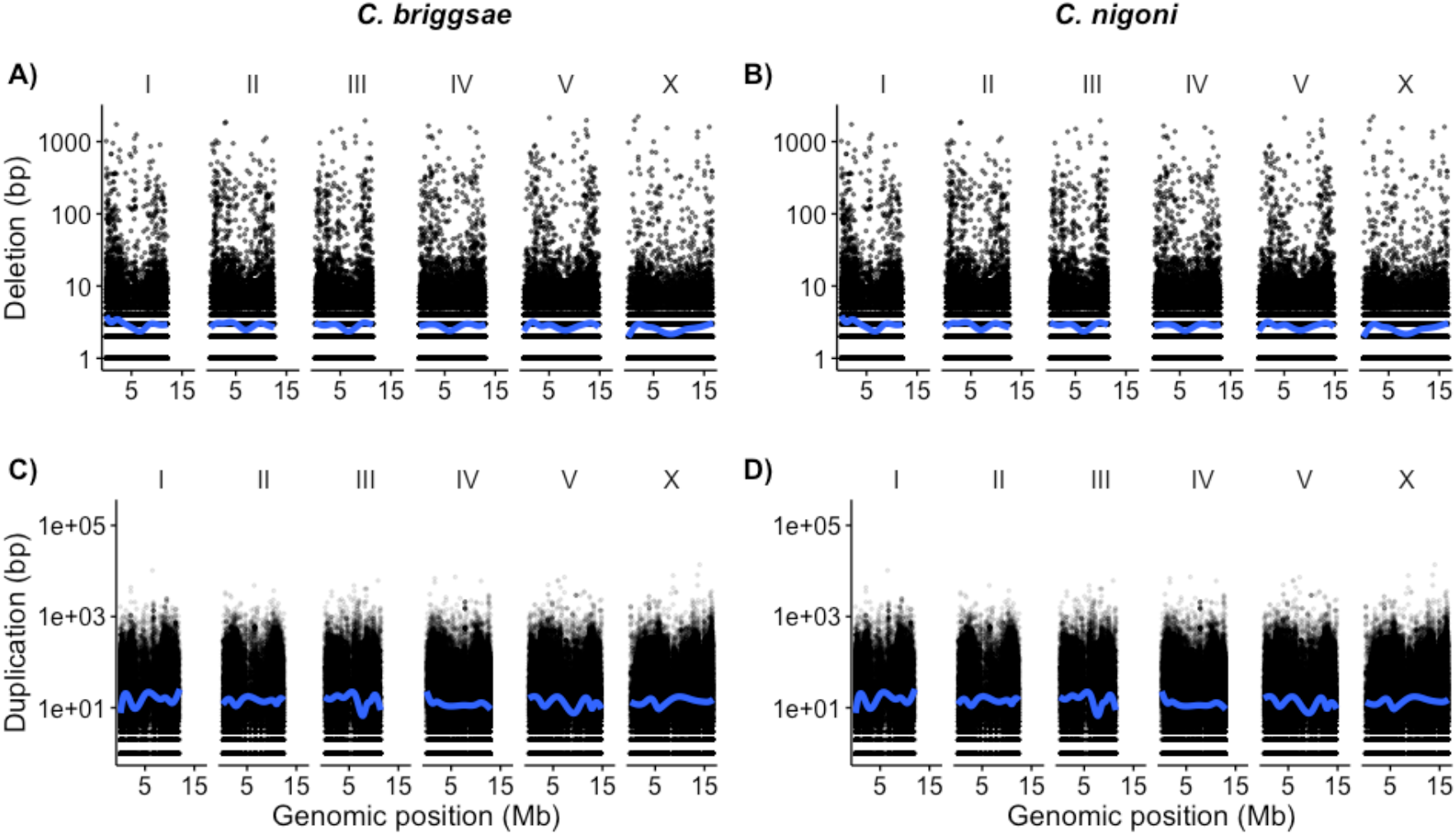
The genomes *C. nigoni*, the most closely related outcrossing species to *C. briggsae*, experienced similar mean and median sizes of Deletions and Duplications

### Transposable element content is not predicted by reproductive mode

The ‘accordion model’ of genome evolution (Kapusta *et al*., 2017) proposes that genomes grow and shrink by a balance of TE-associated expansions and segmental deletions. We find no evidence for this model of genome evolution in the *Caenorhabditis Elegans* group. Despite consistent differences in genome size between outcrossing and self-fertile species, there are no consistent differences in repeat content. Genome size variation within self-fertile species and outcrossing species also does not reflect differences in repeat content. For example, the nematode *C. inopinata* has one of the smallest outcrossing genomes at 122Mb but one of the highest repeat contents at 27.1%. Outcrossing genomes varied between 10.41 and 27.54% repetitive content and self-fertile genomes varied between 9.7 and 21.78% repetitive content (Fig. 1).

It has also been proposed that Class I and Class II TEs might vary predictably with reproductive mode (Boutin *et al*., 2012; Dolgin and Charlesworth, 2006). We find that in the *Caenorhabditis Elegans* group Class II TEs varied similarly across reproductive modes with ranges of 6.96-19.63% in outcrossing genomes and 6.13-16.38% in self-fertile genomes. However, outcrossing genomes had higher proportions of Class I TEs (0.4-4.85%) when compared with self-fertile species (0.68-1%).

To statistically evaluate this Class I TE difference and the ‘accordion model’ of genome size differences via TE expansion we performed a phylogenetic comparative analysis. We mapped the discrete ‘selfer’ or ‘outcrosser’ states onto the phylogeny and distinguished “equal rates”, “symmetrical” and “all rates different” transition matrix models using the small sample size corrected Akaike Information Criteria (AICc). We found the “symmetrical” and “equal” rate transition matrix models both outperformed the “all rates different” model, but were indistinguishable from each other by AICc (Table 1). In all but one case, either the Brownian Motion or the Ornstein-Uhlenbeck global optima model performed best, and although not always distinguishable from each other, always outperformed the Ornstein-Uhlenbeck separate optima model by more than two AICc units (the criteria for significantly different suggested by Burnham and Anderson, 2002).

**Table 1.** Specific TE elements and the AICc scores for a Brownian Motion (BM), single optimum Ornstein-Uhlenbeck model (OU1) and separate self-fertile and outcrossing optima Ornstein-Uhlenbeck model (OU2). Underlined AICc values indicated either the single best model or in some cases the best model and next best model that could not be distinguished by more than 2 AICc units. Eighteen of the 21 TEs were best modeled by a BM process and four of these could not be distinguished from a single optimum OU process. Five TEs were best modeled as a single optima OU process, two of which could not be distinguished from a BM model. Only one of the 21 TEs (the DNA transposon Mutator) was best modeled by a separate optimum OU model. The phylogenetic half-life and stationary variance for Mutator were 0.0011 and 1.35 respectively with greater than 60% of the variance explained by the two optima model (*r*^2^=0.64).

| Element | AICc (BM) | AICc (OU1) | AICc (OU2) |
| --- | --- | --- | --- |
| Repeats(Total) | **<u>73.83</u>** | **<u>74.86</u>** | 77.94 |
| GC % | **<u>24.33</u>** | 28.26 | 27.59 |
| Retroelements | **<u>38.06</u>** | 40.64 | 43.84 |
| Penelope | **<u>-51.02</u>** | -47.09 | -43.06 |
| LINEs | **<u>1.49</u>** | **<u>1.59</u>** | 6.61 |
| L2 CR1 Rex | **<u>-12.83</u>** | **<u>-14.25</u>** | -10.29 |
| R2 R4 NeSL | -37.05 | **<u>-42.28</u>** | -37.49 |
| RTE Bov B | **<u>-17.74</u>** | -13.81 | -9.21 |
| LTR elements | **<u>34.49</u>** | 37.32 | 39.91 |
| BEL Pao | **<u>-19.89</u>** | **<u>-19.86</u>** | -15.10 |
| Gypsy | **<u>20.76</u>** | 24.48 | 27.45 |
| DNA transposons | **<u>57.25</u>** | 61.17 | 64.09 |
| hobo Activator | **<u>-1.73</u>** | 2.20 | 7.03 |
| TC1 IS630 Pogo | **<u>42.08</u>** | 46.01 | 50.93 |
| PiggyBac | **<u>-5.95</u>** | -2.04 | 0.37 |
| Mutator | 51.41 | **<u>51.00</u>** | **<u>49.21</u>** |
| Other (Mirage, P elements, Transib) | 62.05 | **<u>58.63</u>** | 62.87 |
| Helitrons/Rolling Circles | **<u>15.93</u>** | 19.86 | 25.10 |
| Unclassified Repeats | 72.05 | **<u>42.61</u>** | 45.13 |
| Total interspersed repeats | **<u>72.78</u>** | **<u>73.52</u>** | 75.91 |
| Satellites | **<u>12.51</u>** | 16.44 | 21.66 |
| Simple repeats | **<u>-25.85</u>** | -21.92 | -23.05 |

The Ornstein-Uhlenbeck model with separate optima for self-fertile and outcrossing species performed best only for the DNA transposon Mutator and by 1.79 AICc units better than the next best single optima model. The estimates of primary optima for self-fertile species (2.97% +/-0.58) and outcrossing species (6.19% +/-0.44) were greater than 2 standard errors different from each other and given the low level of phylogenetic inertia estimated (*t*_1*/*2_ = 0.11% of the total tree height), were similar to the currently observed mean values within self-fertile and outcrossing species. Mutator elements in self-fertile genomes ranged from 5,234 in *C. briggsae* to 12,076 in *C. tropicalis* JU1373. In comparison, Mutator elements in outcrossing genomes ranged from 24,803 in *C. nigoni* to 47,268 in *C. inopinata*. However, we note that this DNA transposon makes up just 6% and 3% of the respective genomes and cannot explain the much larger difference in overall genome sizes (12-66%) between these two groups.

### Gene family turnover is high across Caenorhabditis

We used orthofinder (Emms and Kelly, 2015) to associate protein-coding genes to 24,574 different orthogroups or gene families. We eliminated orthogroups that were not present in at least half of our *Caenorhabditis* genomes and retained 14,590 gene families. We estimated gene birth and death with the CAFE5 software (Mendes *et al*., 2020; Han *et al*., 2013; Hahn *et al*., 2005) for these gene families. We tested models with 1-9 different birth/death rates (*λ*s) for gene family change and found that a model with 2 birth/death rates (*λ*s) had the highest likelihood. The estimated birth/death rate across the *Caenorhabditis* group (*λ* = 0.4293) was 1-2.5 orders of magnitude higher than that reported in *Drosophila* (Hahn *et al*., 2007) or *Saccharomyces* (Han *et al*., 2013). Rapid gene family expansion and contraction was not limited to one clade or individual species (Fig. 8).

**Fig. 8:**
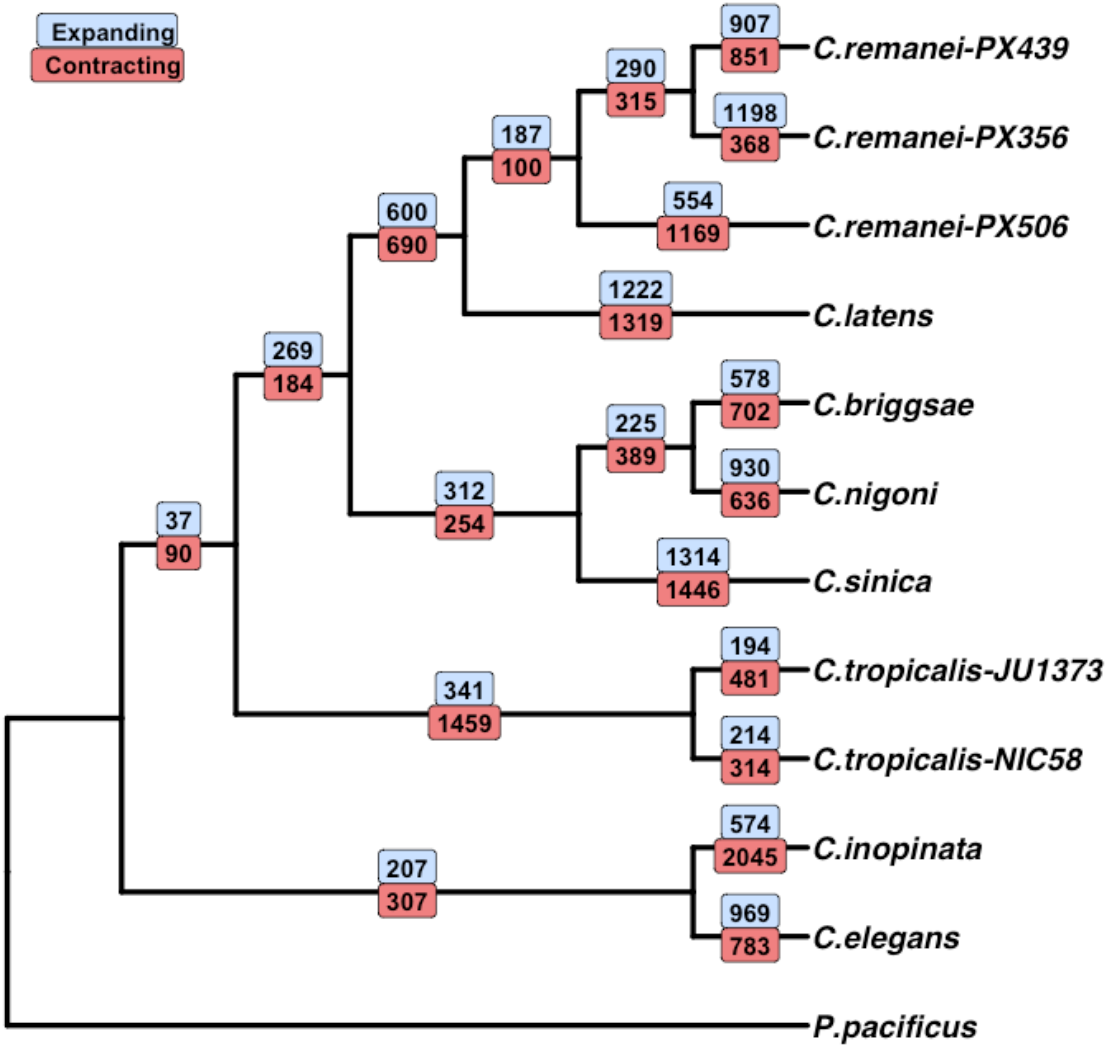
The number of gene family expansions (in blue) and contractions (in red) on each branch of the *Caenorhabditis Elegans* phylogeny.

As an illustration of this we studied the evolution of F-box FBA2 (fbx-a) genes in the *C. elegans*-*C. nigoni* selfing-outcrossing species pair. F-box genes are involved in protein-protein interactions (Kipreos and Pagano, 2000) and implicated in the sexual system in *Caenorhabditis*. For example, recruitment of an F-box gene to a pathway regulating *C. elegans* hermaphrodite development is a crucial piece of evidence for the 3 independent origins of self-fertility in the *Caenorhabditis Elegans* group (Guo *et al*., 2009). F-box genes occur in *Caenorhabditis* genomes in high numbers with at least 377 in *C. elegans* (Wang *et al*., 2021). Of this, 222 are annotated as F-box FBA2 (fbx-a) genes characterized by an F-box domain and an FBA2 domain. The proteins are unevenly spread across the chromosomes with 6 on chromosome IV in *C. elegans* and 92 on chromosome V.

We identified a 2.1kb region of ancestral sequence (Fig. 9) that aligns to multiple regions within a 500kb sections of the *C. elegans* chromosome III, labeled duplicated sequence by ProgressiveCactus (Armstrong *et al*., 2020). The same region of ancestral sequence was not identified in the *C. inopinata* genome and instead aligned poorly to small, disjunct regions across the chromosome. Biological function is not known for F-box FBA genes but they appear to have arisen in high numbers in *Caenorhabditis* through tandem duplications (Wang *et al*., 2021). F-box and F-box FBA genes are abundant in outcrossing *Caenorhabditis* as well, with 1,358 annotated F-box proteins and 412 annotated fbx-a proteins in the outcrossing *C. remanei* PX506. Rapid gene birth and death via small structural mutations were common in both outcrossing and self-fertile in the *Caenorhabditis Elegans* group.

**Fig. 9:**
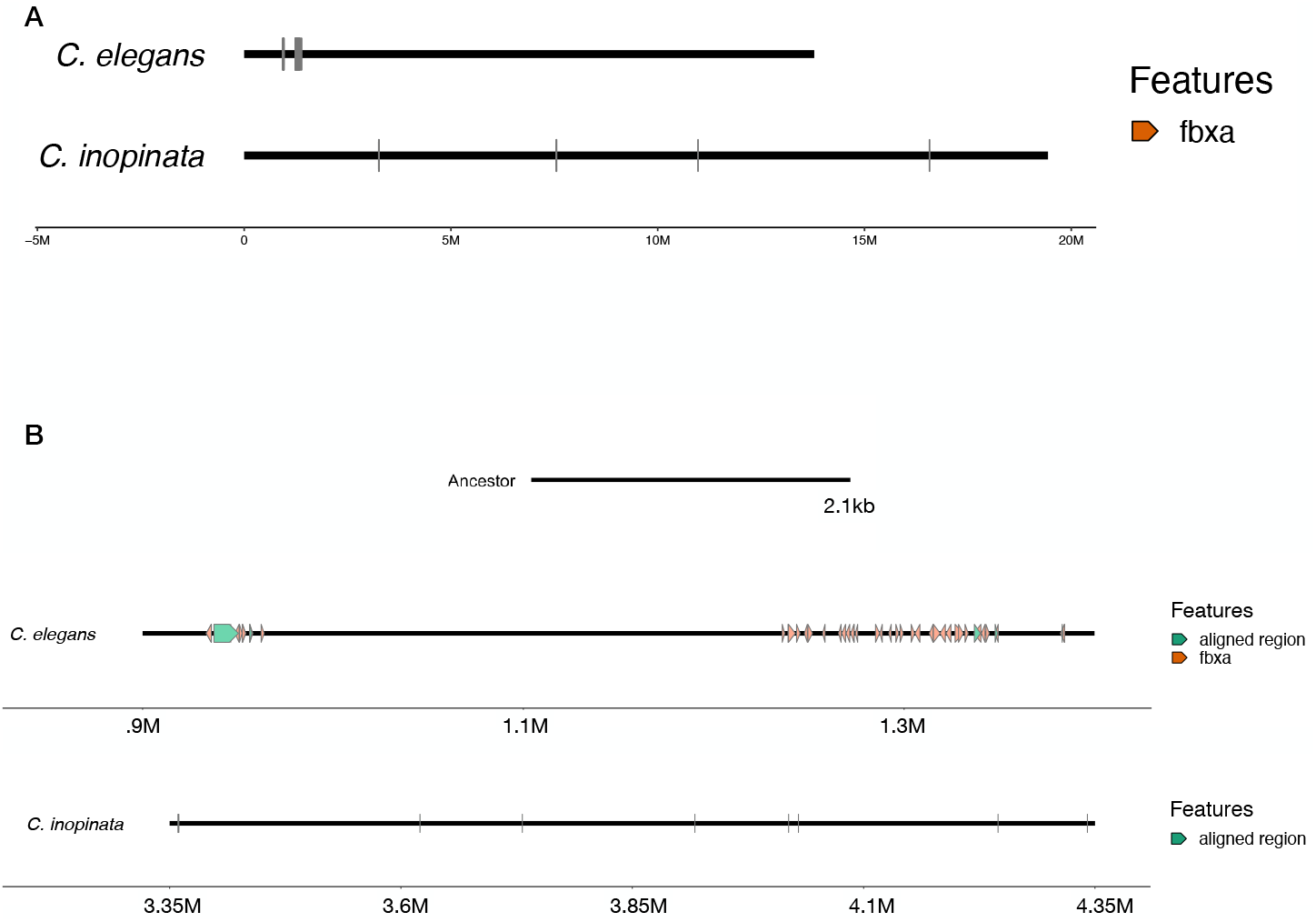
A) A 500kb region of *C. elegans* chromosome III contains 46 annotated fbx-a genes. In comparison there are 4 fbx-a genes spread across 15Mb of chromosome III in *C. inopinata*. B) A 2.1kb region of ancestral sequence aligns to this region multiple times in *C. elegans* with high sequence similarity but aligns weakly to small, disjunct regions in *C. inopinata*.

### Few gene families show parallel changes across self-fertile species

We found that of the 14,590 gene families 595 were expanding or contracting significantly at p*<*0.01. Of these, 71 were decreasing in self-fertile species relative to outcrossing species. We identified an annotated protein domain or *C. elegans* orthologous protein for genes in 30 of these families. Six of these families involved serpentine receptors, chemoreceptors known to be important for *Caenorhabditis* ability to navigate its environment. Three of the families encode F-box associated genes and 10 of the families encode regulatory proteins including transcription factors, DNA polymerase-associated domains, RNA export domains, histone-lysine N-methyltransferases and kinases. We also identified 8 families encoding membrane associated immune and protein responses including peptidases, C-type lectins, ankyrin repeats and actin and chitin associated proteins.

*Caenorhabditis* genomes contain large numbers of each of these protein families and changes in gene number could simply reflect overall changes at the genomic level. We found that F-box associated proteins and kinases were not changing significantly while serpentine receptors (*p* = 0.0142), transcription factors (0.0001) and peptidases were (*p* = 0.0026). Thus, gene family expansions and contractions likely reflect a mix of true selection and drift mediated by high molecular turnover in these species. We did not identify gene families consistently increasing in parallel in selfing species or increasing or decreasing in outcrossing species.

## Discussion

We have presented a fine-scaled analysis of genome evolution across the *Caenorhabditis Elegans* group. Our results contribute to a body of work establishing protein-coding changes as the basis of genome size differences in these worms (Stevens *et al*., 2019; Teterina *et al*., 2020; Thomas *et al*., 2012; Yin *et al*., 2018; Fierst *et al*., 2015; Kanzaki *et al*., 2018; Noble *et al*., 2021). We tested hypotheses proposing different sources of genome size variation between outcrossing and self-fertile species including ‘genome shrinkage’ (Yin *et al*., 2018), the ‘accordion’ model of TE expansion and segmental deletion (Kapusta *et al*., 2017) and differences in Class I and Class II TE dynamics (Boutin *et al*., 2012; Dolgin and Charlesworth, 2006). We found no evidence for these hypotheses as the basis of genome size differences in these species. Instead, our results show genome evolution in *Caenorhabditis* was characterized by high rates of structural variant mutations, particularly insertions and duplications, coupled with rapid nucleotide divergence. Differential genome size in these worms evolved through an excess of duplications and insertions in outcrossing species relative to their self-fertile relatives.

### Caenorhabditis genomes: Duplications, insertions and high rates of gene family evolution

The dominant hypothesis for genome evolution after the advent of self-fertility has been deletion of genes (Fierst *et al*., 2015; Thomas *et al*., 2012; Yin *et al*., 2018; Rodelsperger *et al*., 2018) and loss of DNA (Shimizu and Tsuchimatsu, 2015; Roessler *et al*., 2019; Hu *et al*., 2011). We found no evidence for this in *Caenorhabditis*. Our results suggest instead that *Caenorhabditis* genomes evolved size differences through differential expansion via numerous duplications and insertions coupled with rapid nucleotide divergence. Similar patterns of genome size varying with selfing and outcrossing have been observed in plants including *Arabidopsis* (Hu *et al*., 2011) and *Capsella* (Slotte *et al*., 2013). These comparisons have attempted to test hypotheses regarding mechanisms but reported equivocal results due to rapid evolution, low sample sizes and strong phylogenetic signals.

For example, a large-scale study describing the genomes of 10 newly discovered *Caenorhabditis* reported a substantial phylogenetic influence on genome size with the *Drosophilae* Super-group (the clade at the base of the *Elegans* group) having genome sizes 65-91Mb (Stevens *et al*., 2019) and protein-coding gene counts dipping to just 17,134. Notably, all of these species are outcrossing. The lower bound of these estimates is roughly half that observed for both genome size and gene count in outcrossing *Caenorhabditis* in the *Elegans* group and these two clades comprise the upper and lower bounds for estimates of genome size and gene count across the genus. These results suggest that reproductive transitions and the associated influences on genome evolution in *Caenorhabditis* occur against a background of dynamic genome evolution characterized by frequent SV mutations and genome size changes.

We found that *Caenorhabditis* had high rates of gene family turnover including both birth and death rates, and resulting expansions and contractions across the phylogeny. These rates were 1-2.5 orders of magnitude higher than those previously reported for other metazoans (Hahn *et al*., 2005, 2007; Han *et al*., 2013; Schrader *et al*., 2021). Although the SV mutations are randomly generated, they appear to be contributing to rapid rates of gene duplication and in some instances deletion. These high rates have been experimentally observed in *C. elegans* (Konrad *et al*., 2018; Lipinski *et al*., 2011), to the point that duplications and deletions arise in parallel in replicate populations (Farslow *et al*., 2015).

Given this rich mutational spectrum, it can be challenging to infer which gene families evolved as a consequence of reproductive mode. We found no gene families expanding or contracting consistently in outcrossing species and no gene families were consistently expanding in self-fertile species. Of the 595 gene families that were significantly expanding or contracting across the *Caenorhabditis Elegans* phylogeny, 71 (12%) were contracting in parallel across self-fertile genomes. A smaller protein-coding content in self-fertile species has been repeatedly observed (Fierst *et al*., 2015; Yin *et al*., 2018; Shimizu and Tsuchimatsu, 2015; Roessler *et al*., 2019; Hu *et al*., 2011) and two transcriptome-based studies have reported smaller proteomes and parallel gene family loss in self-fertile *Caenorhabditis* (Thomas *et al*., 2012) and *Pristionchus* (Rodelsperger *et al*., 2018) nematodes. The average gene is 3kb in *Caenorhabditis* and our results show that these reductions contribute little to overall genome size differences. Several gene families were changing in proportion to their overall genome composition and likely reflect genetic drift at the level of SVs.

However, single genes, proteins and protein families can be extremely significant for organismal evolution and our results are not mutually exclusive with important gene losses mediated by the evolution of self-fertility. An important example of this is the male secreted short or *mss* protein family that is required for sperm competition in outcrossing species and has been largely lost in self-fertile *Caenorhabditis* (Yin *et al*., 2018; Yin and Haag, 2019). *Caenorhabditis* have an extremely short mutational path from dioecious outcrossing to self-fertility (Baldi *et al*., 2009), but the transition to androdioecy puts the population into an entirely different selection regime. An outcrossing organism must be able to navigate its environment to find the opposite sex, successfully mate, and frequently deal with many more pathogens and parasites (Andersson, 1994). In multiply-mating species these behavioral and environmental demands exert strong selective pressures on males (and. J Wade, 2003) and, accordingly, male-biased and male-associated genes are preferentially missing in both *Caenorhabditis* (Thomas *et al*., 2012; Yin *et al*., 2018) and *Pristionchus* (Rodelsperger *et al*., 2018). Our results suggest that similar patterns are occurring in proteins responsible for sensory recognition, regulatory systems and membrane associated immune and protein responses. These genomic changes reflect the comprehensive environmental selection pressures that mating systems impose. Thus, while the overall size dynamics were not explained by differential loss of genes or DNA there were still important functional patterns of loss and change influenced by reproductive mode in *Caenorhabditis*.

### Genome size is not predicted by TE content

We found the observed differences in *Caenorhabditis* genome size were not due to TEs. There was no evidence for differences in overall TE content, differential expansion and deletion (the accordion model) (Kapusta *et al*., 2017) or dynamics of Class I/Class II TEs (Boutin *et al*., 2012; Dolgin and Charlesworth, 2006). Our results add to studies showing little change in TE abundance after the evolution of self-fertility (Hu *et al*., 2011; Slotte *et al*., 2013) and little difference in the evolutionary dynamics of Class I and Class II TEs (Nowell *et al*., 2021). However, this absence of evidence is not conclusive evidence against these hypotheses. Models addressing reproductive systems and their influence on TE dynamics make specific predictions based on the relative age of the sexual system and TE invasion into the genome (i.e., recent vs ancient) and are sensitive to variation in parameters like quantitative rates of outcrossing and transposition. These subtle variations may act in populations but not be discernible in broad-scale comparative analyses across multiple types of TEs and long evolutionary divergence times. Further fine-scale studies that are better able to match experimental data with theoretical parameters may result in greater insights into the interactions of reproductive systems and TE evolution.

The one exception to this general lack of differentiation in TEs was the DNA transposon Mutator. Mutator-like elements (MULEs) draw their name from their mutagenic abilities as the TEs are highly active near genes and frequently acquire host gene fragments (Dupeyron *et al*., 2019). Our phylogenetic comparative analysis found the outcrossing primary optimum was roughly twice that of the self-fertile primary optimum for Mutator TEs, which results in two interesting biological questions. First, why is the Mutator content higher in outcrossing species? And second, are Mutator dynamics responsible for the high rates of rearrangements we observed in *Caenorhabditis* genomes?

Mutator transposons were first discovered in maize where it was found that lines with the TE had mutation rates 30x higher than those without (Robertson, 1978). Since then Mutator transposons and Mutator-Like Elements (MULEs) have been extensively studied in maize and used as forward and reverse mutagenesis systems (Lisch, 2015). In addition to high rates of transposition activity Pack-*Mutator* -like transposable elements (Pack-MULEs) modify genes through biased acquisition of GC-rich sequences and preferential insertion near the 5’ end of transcripts (Jiang *et al*., 2011). Mutator-based mutagenesis overwhelmingly affects genes because of these unique sequence-level mechanisms. Multiple generations of self-fertilization can silence Mutator transposons in maize (Robertson, 1986) through DNA methylation (Chandler and Walbot, 1986; Martienssen and Baron, 1994; Slotkin, 2005) that heritably modifies histones (Guo *et al*., 2021).

These results in maize suggest that Mutator transposons may be differentially affected by outcrossing and self-fertilization, and the contribution to mutational dynamics in maize may also occur in nematodes. However, mutator transposons have only recently been discovered in metazoans and there is little known about how their dynamics may be similar to or differ from those observed in plants (Liu and Wessler, 2017). For example, despite the interesting parallel with self-fertilization DNA methylation is absent in *Caenorhabditis* (Simpson *et al*., 1986; Wenzel *et al*., 2011) and any mechanism of Mutator control in worms would have to occur through a separate mechanism. Mutator transposons in nematodes, particularly *Caenorhabditis*, may be an exciting avenue for future studies of mutational dynamics.

### Variation in genome size and protein-coding gene number within conspecific lineages and between interfertile species

Previous studies of genome structure in *Caenorhabditis* have compared between distantly related self-fertile species (Stein *et al*., 2003; Hillier *et al*., 2007), outcrossing species and distantly related self-fertile species (Thomas *et al*., 2012; Fierst *et al*., 2015) or closely related outcrossing-selfing pairs (Kanzaki *et al*., 2018; Yin *et al*., 2018). We sought to evaluate the significance of these comparisons by comparing genome structure and sequence between the conspecifics *C. remanei* PX356, PX439 and PX506 and the interfertile *C. latens*. Similar comparisons have been performed for globally distributed strains of *C. elegans* (Cook *et al*., 2017; Kim *et al*., 2019; Lee *et al*., 2019) and identified a somewhat paradoxical pattern of low nucleotide diversity coupled with high intraspecific tolerance for structural variations to the level of presence-absence variation in entire genes (Lee *et al*., 2022). We find that this pattern is echoed in outcrossing *Caenorhabditis* with variation in genome size, protein-coding gene number and gene family size across strains within the *C. remanei* species.

This genomic structure where a complement of genes is shared among all members of a group and substantial gene presence-absence variation occurs between genomes has been recognized in prokaryotes and termed the “core genome” and “pangenome” (Tettelin *et al*., 2005). The pangenome concept has been applied to a broad range of organisms including fungi, plants and animals (Golicz *et al*., 2020). However, the scale of variation within and between core and pangenomes varies dramatically across these groups. For example in microbes the core genome is frequently *<*10% of the full pangenome reference (van Tonder *et al*., 2014) while in fungi (McCarthy and Fitzpatrick, 2019) and plants (Gao *et al*., 2019) the core genome comprises 80-90% of the pangenome reference. In humans the pangenome fraction is smaller still, comprising an estimated 0.5-1.2% of the reference sequence (Miga and Wang, 2021).

Within *Caenorhabditis* species a core-pangenome structure is difficult to define because many presence-absence variant genes are members of large, diverse protein families (Lee *et al*., 2022). Identifying one-to-one orthologous relationships given an array of highly divergent paralogs within each genome is difficult in *Caenorhabditis* and other groups characterized by rapid molecular changes (Zallot *et al*., 2016). A more useful division for *Caenorhabditis* protein famillies may be those that are sufficient in small number and those that proliferate within individual genomes. For example, the small worms are extremely dependent on external environment and have highly developed repertoires of chemoreceptors with some 10-20% of the total gene complement comprised of chemoreceptors (Robertson and Thomas, 2005; Thomas and Robertson, 2008).

### Holocentric chromosomes and sexually antagonistic selection

In addition to the mutagenic potential of Mutator TEs, there is a cellular mechanism that may contribute to the generation of SV mutations in nematodes. Holocentric chromosomes with diffuse centromeres have been noted in plant and animal species including *C. elegans* (Maddox *et al*., 2004). Holocentrism can stabilize double stranded DNA breaks in meiosis and facilitated gene conversion in the silkworm *Bombyx mori* (Mon *et al*., 2011). High rates of SVs and rearrangements have been noted in other holocentric species including aphids (Mathers *et al*., 2021) and lepidopterans (Hill *et al*., 2019).

One of the most compelling questions to emerge from our analyses is why outcrossing species would experience selection for increased genome size and protein-coding content. Theory suggests that sexual conflict, when two sexes have opposing optima (Rowe *et al*., 1994), can select for the retention and divergence of duplicated genes (Connallon and Clark, 2011). This has been observed in *D. melanogaster* with the tandem duplicate genes *Apollo* and *Artemis* supporting a scenario of duplication and divergence resolving sexually antagonistic selection (VanKuren and Long, 2018). Going from theoretical predictions and single gene studies to broad-scale quantification of the contribution of sexually antagonistic selection to genome size variation across taxonomic groups is challenging. It will require high-quality genome sequences, robust characterizations of protein-coding genes and a developed understanding of the dynamics of sexual and natural selection.

Until recently, the technology used to assemble and characterize genome sequences has had an outsize influence on the inference of genome size, protein-coding genes and TEs and our ability to test theoretical predictions regarding genome evolution. For example, a prominent hypothesis linking large effective population sizes with small genome sizes (Lynch and Conery, 2003) generated predictions and explanatory theory for a pattern *opposite* to that observed in worms, plants and other self-fertile/outcrossing species. As the field of evolutionary biology shifts to newer long read DNA technologies we will be increasingly able to quantify patterns of genome evolution as they apply to phylogenetic groups, reproductive modes and functional systems like chromosome pairing and meiosis. The ambitious Darwin Tree of Life programme at the Wellcome Sanger Institute is aiming to sequence the 70,000 eukaryotic species in the UK and Ireland with long-read technology and permit high-quality, chromosome-scale assemblies (Blaxter *et al*., 2022). These resources will be critical for definitively rejecting and critically evaluating hypotheses regarding genome evolution across eukaryotic life.

### Conclusions

The evolution of genome size is a fundamental question in biology (Gregory, 2005b). Comparative studies across eukaryotes have found that genome size variation across large phylogenetic and physical scales is often explained by repeat content (Elliott and Gregory, 2015; Gregory, 2005a; Kapusta *et al*., 2017). Here, we asked if these dynamics explain genome size variation at smaller physical and phylogenetic scales in species with defined reproductive transitions and correlated genome size differences. We found that smaller scale genome size variation is determined by separate factors. Both outcrossing and self-fertile *Caenorhabditis* experienced numerous gene-associated SV mutations with genomes evolving through duplications, insertions and rapid divergence. TEs and repeats do not explain genome size variation among strains within species, between species or between reproductive modes. Within this landscape of frequent rearrangements we found significant, parallel reductions in gene families across self-fertile species. These encode proteins important for the sensory system, regulatory molecules and membrane-associated immune and protein responses, likely reflecting the global shift in selection pressures created by the transition to self-fertility.

### Materials and methods

### Assembling chromosome-scale sequences for the outcrossing C. remanei PX356 and PX439 and C. latens

We used Oxford Nanopore Technologies to generate DNA libraries for 3 strains of outcrossing *Caenorhabditis*; *C. remanei* PX356, PX439 and *C. latens* PX534 (Sutton *et al*., 2021). The two species are closely related (Felix *et al*., 2014) and partially interfertile (Dey *et al*., 2014). We assembled genome sequences in *<*100 contiguous sequences for each strain and used the contiguous *C. remanei* PX506 assembled sequence (Teterina *et al*., 2020) to scaffold the genome sequences into chromosome-scale pseudomolecules (Supplementary Table 1). Each of the assembled sequences we studied was contained in 6-155 contiguous sequences with the exception of *C. sinica* which is contained in 15,261 sequences. We included *C. sinica* in analyses despite this fragmentation as a representative outcrossing species closer to the root of the *Caenorhabditis Elegans* group. We also included the distantly related *Pristionchus pacificus* to orient and root our phylogenetic analyses.

### Nematode laboratory culture

The *C. remanei* PX356 and PX439 and *C. latens* PX534 strains were graciously provided by the lab of Patrick C. Phillips. Nematodes were cultured on 100mm Nematode Growth Media (NGM) plates seeded with E. coli OP50 (Stiernagle, 2006). For sequencing we collected worms from 5-10 100mm plates by washing with M9 media into 15mL conical tubes. Tubes were placed on a tabletop rocker for 1 hour and then centrifuged to pellet nematodes. To minimize E. coli contamination we removed the M9, added fresh M9, mixed the tubes and pelleted the worms by centrifugation, repeating this process 5 times.

### DNA extraction and sequencing

Our protocol for DNA extraction was described in Sutton et al (2021). Briefly, we froze pelleted worms in liquid nitrogen to rupture cuticles and combined 1.2mL lysis buffer (100mM EDTA, 50mM Tris, 1% SDS) with 20*µ*L Proteinase K (100mg/mL) before incubating at 56C for 30 minutes with shaking. We used phenol chloroform for extraction following Sambrook (2001) and the Short Read Eliminator Kit from Circulomics Inc. (Baltimore, MD) to select high molecular weight DNA.

We used the Oxford Nanopore Technologies (Oxford, UK) SQK-LSK109 ligation sequencing kit for DNA library preparation. Approximately 500-900ng of DNA was sequenced for 48 hours on R9.4.1 RevD fowcells via a gridION X5. We used Guppy v.4.0.11 for basecalling in the ‘-high-accuracy’ mode.

### Genome assembly strategy

We used the Canu v2.0 software package to correct Nanopore libraries (Koren *et al*., 2017). Briefly, Canu’s correction module creates an all-vs-all overlap dataset, uses this to correct individual reads, selects the longest available reads and creates a dataset of user-specified coverage. For the data presented here we used 40x coverage based on an estimated genome size of 130Mb. We assembled the Canu-corrected reads with Flye v.2.8.2 (Kolmogorov *et al*., 2019) and polished the assembled sequences with Pilon v1.23 (Walker *et al*., 2014). We designed this correction, assembly and polishing protocol after extensive simulations and testing (described in Sutton *et al*., 2021). For polishing we used paired-end Illumina libraries previously generated by the laboratory of Patrick C. Phillips and obtained from the NCBI SRA in December 2020 (accessions are listed at the end of this article). We eliminated microbial and other contaminants after polishing with the SIDR software (Fierst and Murdock, 2017). Briefly, SIDR uses ensemble-based machine learning to train a model of sequence identity (i.e., target or contaminant) based on measured predictor variables. Here, the predictors were sequence GC content, read depth of Nanopore libraries aligned to the assembled sequences and *k* -mer frequency distributions with *k* =19.

Residual allelism has been a problem with previous *Caenorhabditis* genome sequences (Barriere *et al*., 2009). We used the purge haplotigs software version 1.1.1 (Roach *et al*., 2018) to identify possible heterozygous regions of the assembled sequences. Briefly, we aligned the ONT libraries to the assembled sequences and produced a read-depth histogram to identify regions of abnormal sequencing depth. The method works under the assumption that alleles will result in ‘split coverage’ with read depth approximately 0.5 that of the homozygous contigs. The read-depth histogram did not have noticeable regions of abnormal coverage and the software identified less than 400,000bp of possible haplotigs in the genome sequences of *C. remanei* PX356, *C. remanei* PX439 and *C. latens* PX534. This represented *<*0.33% of any of the assembled genome sequences and we chose to retain these sequences in the assembled genome because we could not reliably identify them as ‘haplotigs.’

### Creating pseudo-molecules

Previous studies have shown remarkable conservation of large-scale synteny between *Caenorhabditis* (Yin *et al*., 2018; Fierst *et al*., 2015; Stein *et al*., 2003; Teterina *et al*., 2020). We assumed this large-scale synteny is conserved within the interfertile *C. remanei* /*C. latens* species complex and used the chromosome-scale *C. remanei* strain PX506 assembled genome sequence (Teterina *et al*., 2020) to construct pseudo-molecules for *C. remanei* PX356 and PX439 and *C. latens* PX534 with the RagTag software version 2.1.0 (Alonge *et al*., 2021). Briefly, RagTag performs homology-based scaffolding by aligning query sequences to a reference assembled sequence with the minimap2 software (Li, 2018).

### Gene annotation

We annotated protein-coding genes in the assembled sequences of *C. remanei* PX356 and PX439 and *C. latens* PX534 with the BRAKER2 v2.1.6 software (Bruna *et al*., 2021). We used RepeatModeler v2.0.2 (Smit *et al*., 2015) for *de novo* repeat identification and the queryRepeatDatabase.pl script inside RepeatMasker/util to extract Rhabditida repeats (Bao *et al*., 2015). We combined these files to create a library of known and *de novo* repeats and used these with RepeatMasker v4.1.2-p1 (Smit *et al*., 2015) to softmask repeats in the assembled sequence. We aligned RNA-Seq libraries extracted from mixed stage nematode populations to the softmasked sequences with STAR aligner v2.7.9a (Dobin *et al*., 2013). We used this in BRAKER2 with the protein sequences from *C. remanei* PX506 as homology evidence. Only the RNA-Seq libraries were used for training gene predictors. We obtained the assembled genome sequences, protein-coding gene annotations and coding sequence files for the remaining *Caenorhabditis* species from WormBase Parasite (Howe *et al*., 2017) in December 2020 (version WBPS15) with the exception of *C. tropicalis*, which was obtained from the NCBI in February 2021. We performed all of the following analyses (functional annotation, transposable element annotation, whole genome alignment) on the *C. remanei* and *C. latens* assembled genome sequences and protein-coding gene annotations produced by our group and the *Caenorhabditis* assembled genome sequences and protein-coding gene annotations obtained from WormBase Parasite and the NCBI. We used the AGAT suite to calculate gene statistics (Dainat, v080).

### Functional annotation

We used the Interproscan software v5.19 (Finn *et al*., 2017; Zdobnov and Apweiler, 2001) to annotate protein domains and motifs, gene ontologies *et al*., 2016; Kanehisa and Goto, 2000). Briefly, Interproscan searches multiple databases for protein information including PRINTS (Attwood and Beck, 1994; Attwood *et al*., 1994), Pfam (Punta *et al*., 2012), ProDom (Bru *et al*., 2005) and PROSITE (Hulo *et al*., 2005).

### Whole genome alignment

We used ProgressiveCactus version 2.0.4 (Armstrong *et al*., 2020) to align the *Caenorhabditis* genome sequences and measure the spectrum of mutational events. ProgressiveCactus permits reference-free alignment and uses phylogenetic information to estimate parental (ancestral) genome sequences. We used the *Caenorhabditis* phylogeny estimated by Stevens *et al*. (2019) as the basis for the alignment and added the strains *C. remanei* PX356, PX439, PX506 and *C. tropicalis* JU1373 and NIC58 as polytomies. There is some genetic differentiation between strains within each species but the scale of protein divergence at loci conserved across Phylum Nematoda is minimal within strains when compared with the entire phylum (Bird *et al*., 2005). Additionally, the inbreeding process necessary to create homozygous strains can lead to unusual fixations and genetic sampling effects that do not accurately represent the species (Adams *et al*., 2022; Roessler *et al*., 2019). We used *Pristionchus pacificus* as an outgroup to root the alignments. *P. pacificus* is a distant relative of the *Caenorhabditis* group (Dieterich *et al*., 2008) but is the closest relative to the group with a high-quality assembled genome sequence suitable for use as a reference.

The ProgressiveCactus alignment algorithm reconstructs an ancestral sequence for branch points along the phylogeny and mutational events are measured between ancestor and child genomes. ProgressiveCactus defines deletions, insertions, gap deletions and gap insertions by size, where gap deletions and insertions are *<*5bp. Transpositions involve transfer of sequence from one chromosomal region to another while inversions are regions that have reversed orientation. Duplications are sequences that occurred singly in the ancestor and in multiple copies in the child genome.

The output of a ProgressiveCactus (Armstrong *et al*., 2020) alignment is a binary hierarchical alignment (HAL) file (Hickey *et al*., 2013). We used the halSummarizeMutations function to calculate substitutions, transitions, transversions, insertions, (Consortium, 2000) and pathway information (Kanehisa deletions, duplications and transpositions. The halSummarizeMutations function calculates these quantities for each genome relative to the ancestral genome and for estimated ancestral genomes relative to each estimated parent node. We used the halBranchMutations function to create a BED-formatted file with the genomic locations of each of the mutations and the bedtools intersect function to associate these mutations with annotated repeat elements and protein-coding genes (Quinlan and Hall, 2010). We used the hal2fasta function to print the estimated ancestral genome sequences, annotated repeat elements in these with the EDTA software (Ou *et al*., 2019) and associated mutations in ancestral genomes with annotated repeats using bedtools intersect. We could not reconstruct protein-coding genes in estimated ancestral sequences because the accuracy of protein-coding gene annotation is dependent on nucleotide-level features like start codons and accurate intron-exon boundaries that are not prioritized in Progressive Cactus alignment. However, this approach did allow us to analyze insertions and deletions in repeat elements and specific TE families.

### TE annotation

We used the Extensive *de novo* TE Annotator software version 2.0 (EDTA; Ou *et al*., 2019) to identify repeats in *Caenorhabditis* genome sequences. EDTA uses a number of different open-source tools to identify TE candidates in genome sequences and combines these with annotated repeats using known coding sequences (CDS) to eliminate false positive TEs. We used the Rhabditida repeats extracted from RepeatMasker (Bao *et al*., 2015) and the CDS obtained from BRAKER2 (Bruna *et al*., 2021) annotation, WormBase Parasite (Howe *et al*., 2017) and the NCBI. Long terminal repeat retrotransposons (LTRs) were identified with LTR Harvest (Ellinghaus *et al*., 2008), LTR Finder (Xu and Wang, 2007), LTR retriever (Ou and Jiang, 2018, 2019) and Generic Repeat Finder (Shi and Liang, 2019). Generic Repeat Finder (Shi and Liang, 2019) was also used to identify Terminal Direct Repeats (TDRs), Miniature Inverted repeat Transposable Elements (MITEs) and Terminal Inverted Repeats (TIRs). TIRs were also identified with TIR-Learner (Su *et al*., 2019). Helitron transposons were identified with HelitronScanner (Xiong *et al*., 2014) and TEsorter used to classify identified TEs (Zhang *et al*., 2019). EDTA (Ou *et al*., 2019) calculates the number, base pairs and percentage of the genome covered by different TEs but a large proportion of the TIR (DNA) elements were not assigned to families found in the Sequence Ontology database and classified as generic ‘repeat region.’ To accurately characterize these we used the RepeatMasker (Bao *et al*., 2015) annotation table with the EDTA-identified TEs to calculate the number, base pairs and percentage of the genome covered by different classes and types of TEs.

### Phylogenetic comparative analyses

Hansen (1997) introduced the use of an Ornstein-Uhlenbeck model to test hypotheses of trait adaptation to different niches mapped on a phylogeny. The Ornstein-Uhlenbeck process includes parameters that capture deterministic movement of species trait values toward optimal states that can vary as a function of environmental or ecological variables. Hansen (1997) termed these states ”primary optima” defined as the average expected trait values (the local optima) for many species adapting to a given primary niche. The idea is that ”secondary” selective factors average out across species, leaving the common effect of the primary niche on trait values. Hypotheses about adaptation can be tested by estimating the primary optima for different states of the environmental or ecological variables and asking if they differ as predicted by the hypotheses (see Hansen (2014) for a detailed argument).

Mathematically, a simple Ornstein-Uhlenbeck process is described by the stochastic differential equation:

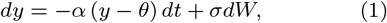

where *dy* is the change in a given species’ mean trait value, *y*, over a short time interval *dt, θ* is the primary optimum, *α* determines the rate of adaptation toward the primary optimum, *dW* represents independent normally distributed stochastic changes with mean zero and unit variance over a unit of time, and *σ* is the standard deviation of these changes. The *σ* parameter is more readily interpretable when expressed as a stationary variance of the process, *v* = *σ*^2^*/*2*α*, which is the variance among species within a niche after a long period of independent evolution. Using SLOUCH (Hansen *et al*., 2008; Kopperud *et al*., 2018), one of many R packages that implement these methods, we investigated whether nematodes sharing a given niche (here, dioecious or androdioecious reproduction) tend to be more similar compared to taxa sharing a different niche in terms of their TE compositions whilst simultaneously estimating and controlling for the levels of adaptation and phylogenetic inertia in the clade (see Hansen, 2014, Mahler and Ingram, 2014 and O’Meara and Beaulieu, 2014 for general reviews). We analyzed differences in total repeat content and number for 21 different classes of repeats including retroelements like SINES, LINES and LTR elements, DNA transposons like PiggyBac and Mutator, rolling circle/Helitrons, unclassified elements and simple repeats (Table 1).

We mapped ‘selfer’ and ‘outcrosser’ as discrete states onto the phylogeny using maximum likelihood employed by the ace function in the Ape R package (Paradis and Schliep, 2018). The ancestral states along each branch were discretized by choosing the state with the highest probability on each branch for the best model. We used SLOUCH (Hansen *et al*., 2008; Kopperud *et al*., 2018) to fit a Brownian motion model and two Ornstein-Uhlenbeck models, one with a single global optimum and one with separate primary optima for selfers and outcrossers, for each of the twenty-two TEs studied here. The small sample size corrected Akaike Information Criteria (AICc) was used to determine which of the three models best captured TE evolution.

### Orthology assignment

We used OrthoFinder version 2.5.4 (Emms and Kelly, 2015) to identify orthologous and paralogous genes in our *Caenorhabditis* genomes. We selected the longest isoform for each gene with the OrthoFinder primary transcript.py. Briefly, OrthoFinder aligns proteomes with DIAMOND (Buchfink *et al*., 2015, 2021) and uses a Markov Cluster Algorithm to assign proteins to orthogroups.

### Gene birth and death analyses

We used the orthogroups assigned by OrthoFinder (Emms and Kelly, 2015) to calculate gene family expansions and contractions with the CAFE5 software (Mendes *et al*., 2020). CAFE5 assumes that each gene family has at least one representative at the base of the tree and we eliminated the distantly related *P. pacificus* from the analysis. *C. sinica* has the most fragmented assembled sequence of the species we studied here but its annotated protein-coding gene complement is large enough to fit this requirement and we included it throughout this manuscript to orient our analyses and reduce bias that might be introduced by overly distant species like *P. pacificus*, assembled sequences with retained allelism like the outcrossing *C. brenneri* (Barriere *et al*., 2009) or possible reduced complements of protein-coding genes like the self-fertile *C. elegans* (Stevens *et al*., 2019). We used phytools (Revell, 2012) to create a dichotomous, ultrametric phylogenetic tree from the Stevens *et al*. (2019) estimated phylogeny.

We estimated an error model with the CAFE5 ‘-e’ option and used this error model in further analyses. CAFE5 uses maximum-likelihood estimation to fit gene birth-death rate parameters or *λ* values based on a user specifying the number of discrete *λ* values. Gene families changing significantly are those experiencing rapid expansions or contractions across segments of the phylogenetic tree. We fit models with 2-8 *λ* values and found that the model with *λ*=2 had the highest likelihood given the dataset.

To focus on gene families that were expanding or contracting in parallel in selfing and outcrossing lineages we extracted orthogroups that were identified as significant (*p <* 0.01) in the CAFE5 analyses and implemented a set of Boolean rules. If an orthogroup was stable or decreasing in all selfing lineages and stable or increasing in all outcrossing lineages we defined it as ‘Decreasing in Selfers.’ Similarly, if an orthogroup was stable or decreasing in all outcrossing lineages and stable or increasing in all selfing lineages we defined it as ‘Decreasing in Outcrossers.’ If an orthogroup was stable or increasing in all selfing lineages and stable or decreasing in all outcrossing lineages we defined it as ‘Increasing in Selfers.’ Similarly, an orthogroup that was stable or decreasing in all outcrossing lineages and stable or increasing in all selfing lineages was ‘Decreasing in Outcrossers.’ We searched the functional annotations for information on any member of these orthogroups and searched *C. elegans* for orthologous genes as these may also provide functional information.

## Accessions

*C. remanei* PX356 Bioproject PRJNA248909

*C. remanei* PX439 Bioproject PRJNA248911

*C. remanei* PX506 Bioproject PRJNA577507

*C. latens* PX534 Bioproject PRJNA248912

*C. briggsae* AF16 Bioproject PRJNA20855

*C. nigoni* JU1422 Bioproject PRJNA384657

*C. elegans* N2 Bioproject PRJNA158, PRJNA13758

*C. tropicalis* NIC58, JU1373 Bioproject PRJNA662844

*C. inopinata* NKZ35 Bioproject PRJDB5687

*C. sinica* ZZY0401 Bioproject PRJNA194557 Bioinformatic scripts, software and workflows associated with this project are located at https://github.com/jannafierst/Worm-nomics.

## Supplementary Material

Supplementary materials are available at XXXX.

## Acknowledgments

The authors would like to thank Rohit Kapila, Victoria (Tori) Eggers, Justin Rosario, Ana Perez-Sanchez, Jessica Gonzalez and Adam Trautwig for helpful comments and feedback. PEA and JMS were supported by National Alumni Association Fellowship through the University of Alabama Alumni Association. JP was supported by the US National Science Foundation award DEB 2225683 and Florida International University start-up funds. JLF was supported by the US National Science Foundation awards EF 1921585 and DEB 1941854 and Florida International University start-up funds.

## Notes

https://github.com/jannafierst/Worm-nomics

### Competing Interest Statement

The authors have declared no competing interest.

### Summary of Updates

Author order updated on BioRXiv submission to reflect .pdf and manuscript.

