## Supplemental Tables for "Genome size changes by duplication, insertion and divergence in Caenorhabditis worms"

*Journal Title Here*, XXXX, 1–1
doi: DOI HERE
Advance Access Publication Date: Day Month Year
Paper

### Genome size changes by duplication, insertion and divergence in Caenorhabditis worms

**Joshua D. Millwood,[1] John M. Sutton,[1,2] Paula E. Adams,[1,3] Jason Pienaar[4,5,7] and Janna L. Fierst[6,7]***

[1]Department of Biological Sciences, The University of Alabama, Tuscaloosa, AL 35487, [2]Current address: Absci, Vancouver, WA 98663, [3]Current address: Department of Biological Sciences, Auburn University, Auburn, AL 36830, [4]Institute of the Environment, [5]Center for Tropical Botany, [6]Biomolecular Sciences Institute and [7]Department of Biological Sciences, Florida International University, Miami, FL 33199

FOR PUBLISHER ONLY Received on Date Month Year; revised on Date Month Year; accepted on Date Month Year

#### Abstract

**Table 1.** Assembly statistics for the *Caenorhabditis* species and strains. Assembled sequence and contig # are raw assembly output and pseudomolecule contig # are after homology-based scaffolding using the *C. remanei* PX506 sequence as a reference. The *C. remanei* PX506 pseudomolecules were generated with Hi-C technology.

| | Assembled sequence (Mb) | Contig # | Pseudomolecule sequence (Mb) | Scaffold # |
|---|---|---|---|---|
| *C. remanei* PX506 | 130.48 | 197 | 124.80 | 6 |
| *C. remanei* PX356 | 124.50 | 70 | 124.07 | 17 |
| *C. remanei* PX439 | 132.05 | 53 | 132.00 | 17 |
| *C. latens* | 120.37 | 97 | 119.66 | 38 |
| *C. briggsae* | 105.42 | 12 | | |
| *C. elegans* | 100.81 | 7 | | |
| *C. tropicalis* NIC58 | 81.32 | 7 | | |
| *C. tropicalis* JU1373 | 80.98 | 44 | | |
| *C. inopinata* | 122.58 | 6 | | |
| *C. nigoni* | 129.44 | 155 | | |
| *C. sinica* | 130.39 | 15,261 | | |

© XXXX XXXX. 

**Table 2.** Insertion and Inversion/Transposition sizes for *Caenorhabditis* species and strains with assembled sequences contained in <100 contiguous pieces. Insertions and Inversions/Transpositions are calculated in the species genome coordinates.

| | Insertion number (genes) | Mean size (Median) +/- sd | Insertion number (TEs) | Mean size (Median) +/- sd | Inversion/Transposition number (genes) | Mean size (Median) +/- sd | Inversion/Transposition number (TEs) | Mean size (Median) +/- sd |
|---|---|---|---|---|---|---|---|---|
| *C. remanei* PX506 | 52906 | 38 (3) +/- 166 | 49571 | 186 (7) +/- 498 | 1171 | 7148 (484) +/- 15121 | 1184 | 2672 (362) +/- 5996 |
| *C. remanei* PX356 | 80509 | 30 (3) +/- 132 | 46695 | 140 (6 ) +/- 456 | 1046 | 2107 (265) +/- 6367 | 1170 | 2555 (593) +/- 6130 |
| *C. remanei* PX439 | 76506 | 34 (3) +/- 162 | 44695 | 120 (5) +/- 398 | 1113 | 2083 (239) +/- 5489 | 1286 | 3009 (687) +/- 5909 |
| *C. latens* | 176044 | 29 (4) +/- 153 | 64298 | 139 (7) +/- 435 | 1000 | 260 (55) +/- 771 | 619 | 358 (88) +/- 734 |
| *C. briggsae* | 242671 | 33 (3) +/- 169 | 58192 | 108 (5) +/- 372 | 1028 | 295 (92) +/- 581 | 420 | 386 (126) +/- 862 |
| *C. elegans* | 354081 | 78 (5) +/- 292 | 35386 | 385 (10) +/- 840 | 958 | 117 (29) +/- 250 | 158 | 132 (48) +/- 255 |
| *C. tropicalis* NIC58 | 18988 | 27.36 (3) +/- 278 | 4679 | 661 (4) +/- 2307 | 551 | 56999 (845) +/- 103358 | 140 | 13000 (260) +/- 42490 |
| *C. inopinata* | 224273 | 149 (5) +/- 536 | 90250 | 903 (322) +/- 1421 | 799 | 165 (62) +/- 334 | 501 | 161 (74) +/- 330 |
| *C. nigoni* | 214105 | 49 (3) +/- 496 | 62558 | 207 (5) +/- 1257 | 1284 | 171 (49) +/- 368 | 577 | 200 (87) +/- 330 |

**Table 3.** Deletion and Duplication sizes for *Caenorhabditis* species and strains with assembled sequences contained in <100 contiguous fragments. Deletions and Duplications are calculated in ancestor genome coordinates and can not be associated with annotated gene or TE features.

| | Deletion number | Mean size (Median) +/- sd | Duplication number | Mean size +/- sd |
|---|---|---|---|---|
| *C. remanei* PX506 | 61380 | 6 (2) +/- 14 | 211249 | 85 (21) +/- 238 |
| *C. remanei* PX356 | 154383 | 9 (2) +/- 41 | 140127 | 83 (22) +/- 217 |
| *C. remanei* PX439 | 128950 | 9 (2) +/- 38 | 179330 | 88 (22) +/- 219 |
| *C. latens* | 85313 | 4 (2) +/- 9 | 304085 | 61 (17) +/- 165 |
| *C. briggsae* | 201676 | 4 (2) +/- 16 | 126817 | 67 (16) +/- 238 |
| *C. elegans* | 193488 | 4 (3) +/- 10 | 346466 | 32 (9) +/- 98 |
| *C. tropicalis* NIC58 | 4052 | 6 (2) +/- 47 | 22736 | 104 (24) +/- 353 |
| *C. inopinata* | 141412 | 4 (3) +/- 18 | 341310 | 46 (11) +/- 123 |
| *C. nigoni* | 71457 | 8 (3) +/- 49 | 551878 | 38 (11) +/- 108 |
